# Berberine is a novel mitochondrial calcium uniporter (MCU) inhibitor that disrupts MCU-EMRE assembly

**DOI:** 10.1101/2024.08.18.607892

**Authors:** Haixin Zhao, Siqi Chen, Nian Cao, Wenjun Wu, Guangqin Liu, Jun Gao, Jiayi Chen, Ting Li, Dingyi Lu, Lingmin Zeng, Haizhen Zhu, Weina Zhang, Qing Xia, Teng Li, Tao Zhou, Xue-Min Zhang, Ai-Ling Li, Xin Pan

## Abstract

The mitochondrial calcium uniporter (MCU) complex mediates Ca^2+^ entry into mitochondrial, which plays a crucial role in regulating cellular energy metabolism and apoptosis. Dysregulation of MCU is implicated in various diseases, such as neurodegenerative disorders, cardiac diseases and cancer. Despite its importance, developing specific and clinically viable MCU inhibitors has been challenging. Here, we identify Berberine, a well-established drug with a documented safety profile, as a potent MCU inhibitor through a virtual screening of an FDA-approved drug library. Berberine localizes within mitochondria and directly binds to the juxtamembrane loop domain of MCU. This binding disrupts the interaction of MCU with its essential regulator, EMRE, thereby inhibiting rapid Ca^2+^ entry into the mitochondria. Notably, Berberine pretreatment reduces mitochondrial Ca^2+^ overload and mitigate ischemia/reperfusion-induced myocardial injury in mice. Our findings establish Berberine as a potent MCU inhibitor, offering a safe therapeutic strategy for diseases associated with dysregulated mitochondrial calcium homeostasis.

## Introduction

The entry of Ca^2+^ ions into mitochondria primarily occurs through a highly selective Ca^2+^ channels known as mitochondrial Ca^2+^ uniporter (MCU) (*1*). This process is crucial for regulating various cellular processes, including mitochondrial dynamics, energy metabolism and cell death (*2*). The MCU complex, along with its regulatory components MICU1, MICU2, and EMRE, ensures precise control of mitochondrial Ca^2+^ uptake, which is crucial for maintaining cellular homeostasis (*3*). However, under pathological conditions, dysregulation of mitochondrial Ca^2+^ homeostasis may lead to detrimental effects. Excessive Ca^2+^ accumulation in mitochondria, referred to as mitochondrial Ca^2+^ overload, triggers mitochondrial permeability transition pore opening, leading to mitochondrial membrane depolarization, release of pro-apoptotic factors, and ultimately leading to cell death. These processes contribute to various diseases, including neurodegenerative disorders, cardiac diseases, and ischemia/reperfusion injury (*4, 5*).

Despite the critical role of MCU in disease pathophysiology, developing effective therapeutic interventions targeting mitochondrial Ca^2+^ regulation remains challenging. Existing MCU inhibitors, such as ruthenium red and Ru360, exhibit poor delivery and toxicity issues (*6, 7*). More recent inhibitors, including DS16570511 and mitoxantrone, have shown promise in preclinical studies but are hampered by toxicity, limiting their clinical applicability (*8, 9*). Ideal MCU inhibitors that is suitable for clinical application must meet several criteria: 1) efficient penetration of cellular and mitochondrial membranes to effectively inhibit mitochondrial Ca^2+^ uptake; 2) selective modulatation of mitochondrial Ca^2+^ uptake without affecting cytosolic Ca^2+^ signaling, ensuring normal cellular functions; 3) preservation of mitochondrial membrane potential, which is vital for mitochondrial function; and 4) low toxicity at therapeutically relevant concentrations to ensure safety and minimal adverse effects.

In response to the demand for safe and effective MCU inhibitors, we employed virtual molecular docking followed by a functional screening of an FDA-approved drug library. This led to the identification of Berberine, a natural alkaloid with established safety profiles, as a novel MCU inhibitor. Berberine localizes to mitochondria, directly binds to MCU, and disrupts the assembly of the MCU-EMRE complex. Importantly, Berberine significantly reduces mitochondrial Ca^2+^ overload, providing cardioprotection against ischemia/reperfusion-induced myocardial injury in mice. This discovery positions Berberine as a promising candidate for further development in the broader context of diseases involving mitochondrial Ca^2+^ dysregulation.

## Results

### Screening for potential MCU inhibitors from an FDA-approved drug library

MCU functions as a major gate of Ca^2+^ flux from cytosol into the mitochondrial matrix. Targeting MCU may serve as therapeutic strategies for many diseases, such as pathological cardiac cell death (*10–12*). To closely emulate MCU-targeted clinical treatment, we employed a screening approach using an FDA-approved drug library comprising 2,816 compounds (Fig. 1A). In our initial step, we performed a virtual screening via MCU structure-based molecular docking and identified 120 hits with a docking score of < −7.5 kcal/mol, suggesting their potential MCU binding capacity (fig. S1A). To assess the impact of these hits on mitochondrial Ca^2+^ uptake, we established HeLa cells expressing 4mt-GCaMP6, a fluorescent indicator reflecting mitochondrial Ca^2+^ levels (*13*). Subsequently, we generated a screening system for mitochondrial Ca^2+^ uptake activity using permeabilized Hela cells. Exposure to 100 μM CaCl_2_ led to a rapid increase of the fluorescent signal, indicating mitochondrial Ca^2+^ uptake (fig. S1B). Ru360, a well-known MCU inhibitor (*6, 7*), effectively blocked Ca^2+^ entry into mitochondria (fig. S1, B and C). In a small-scale screen using the 120 hits, four small molecules demonstrated at least 50% inhibition of mitochondrial Ca^2+^ uptake at a concentration of 10 µM (fig. S1, B to D). Among them, Berberine exhibited the highest inhibitory effect, reaching up to 87% (fig. S1D). To rule out the possibility that the observed effect was a secondary consequence of mitochondrial membrane potential disruption, the major driving force for Ca^2+^ uptake within the organelle (*14, 15*), we performed TMRM staining analysis. The mitochondrial decoupler, CCCP, induced a decrease in TMRM intensity. All tested drugs, except for Berberine, including Trifarotene, Eltrombopag, and Tipranavir, significantly decreased the mitochondrial membrane potential (fig. S1E). Therefore, this screening process identified Berberine, a natural compound derived from medicinal herbs, as a promising candidate for further investigation as MCU inhibitor.

**Fig. 1.**
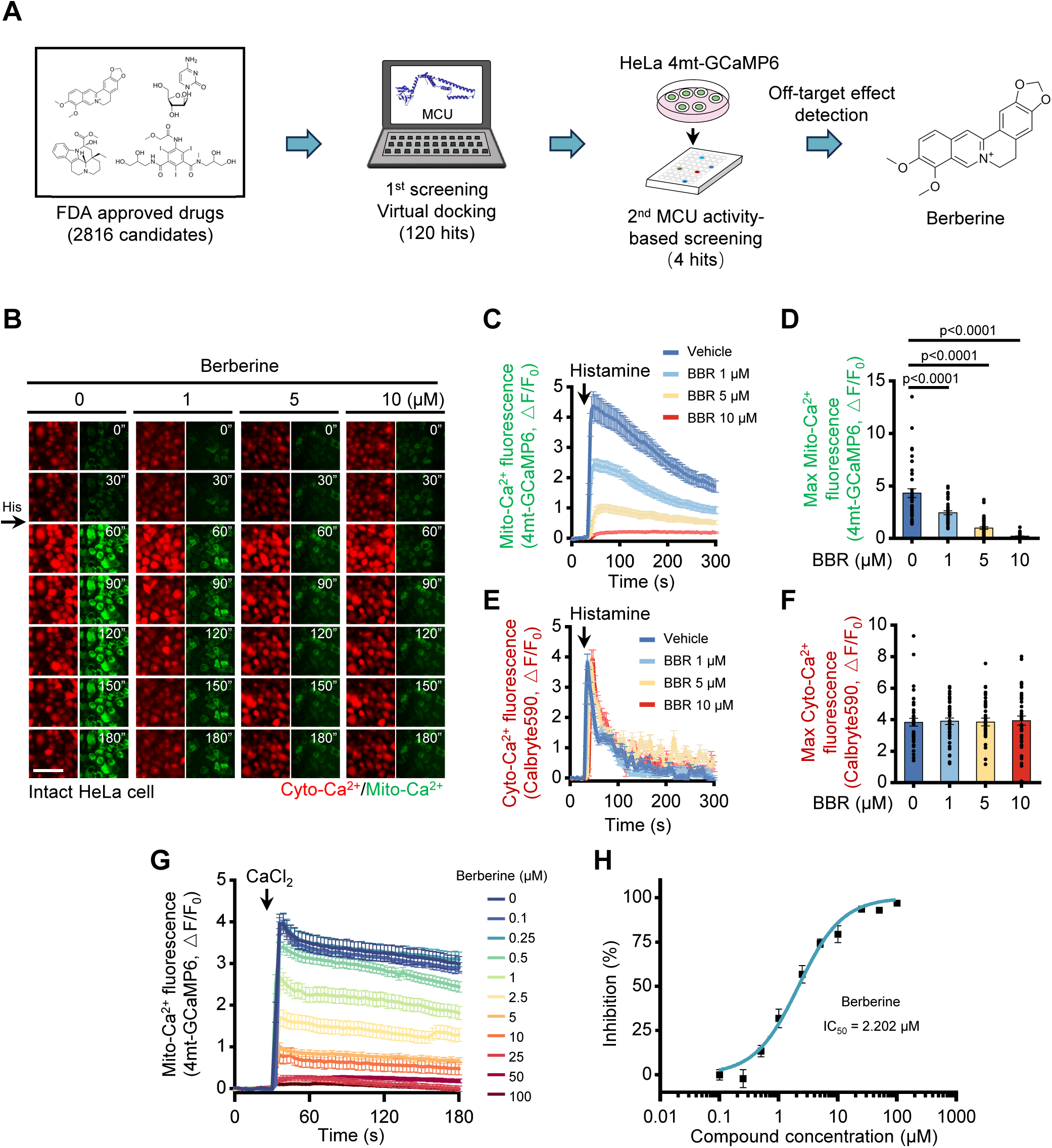
The discovery of Berberine as a novel MCU inhibitor. (A) Schematic of screening workflow. (B) Snapshot of cytosolic (left, red) and mitochondrial (right, green) Ca^2+^ transient images from time-lapse movies of representative HeLa cells stained with Calbryte590 and 4mtGCaMP6 following 10 μM Histamine (His) stimulation. Cells were pretreated with various concentrations of Berberine. The time displayed on the images is in seconds. Scale bar, 50 μm. (C) Representative traces of mitochondrial Ca^2+^ level (indicated by 4mtGCaMP6) in HeLa cells stimulated with 10 μM Histamine and pretreated with different concentrations of Berberine. Data are shown as mean ± s.e.m. n = 40 cells for each condition. (D) Measurement of the maximal amplitudes of the mitochondrial Ca^2+^ traces in (C). Data are shown as the mean ± s.e.m. n = 40 cells for each condition. *P* value was analyzed using one-way ANOVA. (E) Representative traces of cytosolic Ca^2+^ level (indicated by Calbryte590) in HeLa cells stimulated with 10 μM Histamine and pretreated with different concentrations of Berberine. Data are shown as the mean ± s.e.m. n = 40 cells for each condition. (F) Measurement of the maximal amplitudes of the mitochondrial Ca^2+^ traces in E. Data are shown as the mean ± s.e.m. n = 40 cells for each condition. *P* value was analyzed using one-way ANOVA. (G) Traces of mitochondrial Ca^2+^ level (indicated by fluorescence of 4mt-GCaMP6) in digitonin-permeabilized HeLa cells upon the addition of Ca^2+^ under varying concentrations of Berberine. (H) Dose-dependent inhibition of mitochondrial Ca^2+^ uptake activity by Berberine with an IC_50_ of 2.202 μM.

### Berberine inhibits mitochondrial Ca^2+^ uptake

To assess Berberine’s ability to traverse cellular membranes and inhibit mitochondrial Ca^2+^ uptake, we evaluated its inhibitory effects in intact cells by initiating intracellular Ca^2+^ signaling with histamine stimulation while simultaneously monitoring cytosolic and mitochondrial Ca^2+^ dynamics (Fig. 1B). Berberine specifically inhibited mitochondrial Ca^2+^ uptake, as indicated by the green fluorescent indicator 4mt-GCaMP6 (Fig. 1, C and D), without affecting the histamine-induced cytosolic Ca^2+^ signals, as evidenced by the red fluorescent dye Calbryte590 (Fig. 1, E and F). Similar results were obtained using other Ca^2+^ indicators, cyto-GCaMP6, or 4mt-GCaMP8 in HeLa cells (fig. S2, A to D). These findings suggest that berberine can freely traverse the cellular membrane, thereby inhibiting mitochondrial Ca^2+^ uptake without impacting the cytosolic Ca^2+^ signals.

Next, to further quantify berberine’s inhibitory effect on mitochondrial Ca^2+^ uptake, we permeabilized the cells with digitonin, allowing equal doses of calcium ions to be added outside the mitochondrial membrane, thus evaluating the capacity of mitochondria to absorb Ca^2+^ at different concentrations of Berberine. Berberine exhibited a potent inhibition of mitochondrial Ca^2+^ uptake in a dose-dependent manner (Fig. 1G), with an IC_50_ value determined to be 2.202 µM (Fig. 1H).

To evaluate the safety of Berberine in cells, we treated the cells with increasing amount of Berberine, and confirmed that up to 10 µM Berberine treatment did not affect mitochondrial membrane potential by TMRM staining (fig. S2, E and F). We further investigated the cytotoxicity of Berberine to understand its potential impact on cell growth. It showed that at lower doses, specifically between 5-10 µM, which are sufficient to inhibit MCU, Berberine exhibited minimal cytotoxic effects (fig. S2G). This indicates that Berberine is an effective MCU inhibitor with low cytotoxicity, making it a relative safe option at these concentrations.

### Direct binding of Berberine to MCU

Molecular docking screenings indicate that BBR may directly bind to the MCU monomer (Fig. 1A, and fig. S1A). This leads us to suggest that Berberine possesses the unique ability to directly traverse cellular membranes, gaining entry into mitochondria to target and subsequently inhibit MCU. Previous studies have documented the subcellular localization of Berberine within mitochondria (*16, 17*). Exploiting Berberine’s autofluorescence property, we also observed its cellular penetration at 10 μM, co-localizing with MitoTracker stain (fig. S3, A and B), reinforcing our hypothesis.

To further validate our hypothesis, we synthesized biotin-conjugated Berberine (Biotin-BBR) (Fig. 3A). The Biotin-BBR exhibited similar mitochondrial localization (Fig. 2, B and C) and inhibition efficacy as unmodified Berberine (fig. S3, C and D), suggesting that the Biotin-BBR retains the ability of interacting with the same molecular targets as the unmodified Berberine. Consistent with these results, Biotin pull-down assays demonstrated that Biotin-BBR could interact with both exogenous and endogenous MCU in cell lysate (Fig. 2D, and fig. S3E). *In vitro* pull-down assays further confirmed that Biotin-BBR directly binds to recombinant glutathione-S-transferase (GST)-MCU (Fig. 2E). Notably, this binding was diminished when unlabeled Berberine competitively interacted with GST-MCU, indicating a consistent binding site and confirming the specificity of the binding (Fig. 2E). Microscale thermophoresis (MST) analysis further confirmed that the affinity between GST-MCU and Berberine was ∼8 μM (Fig. 2F).

**Fig. 2.**
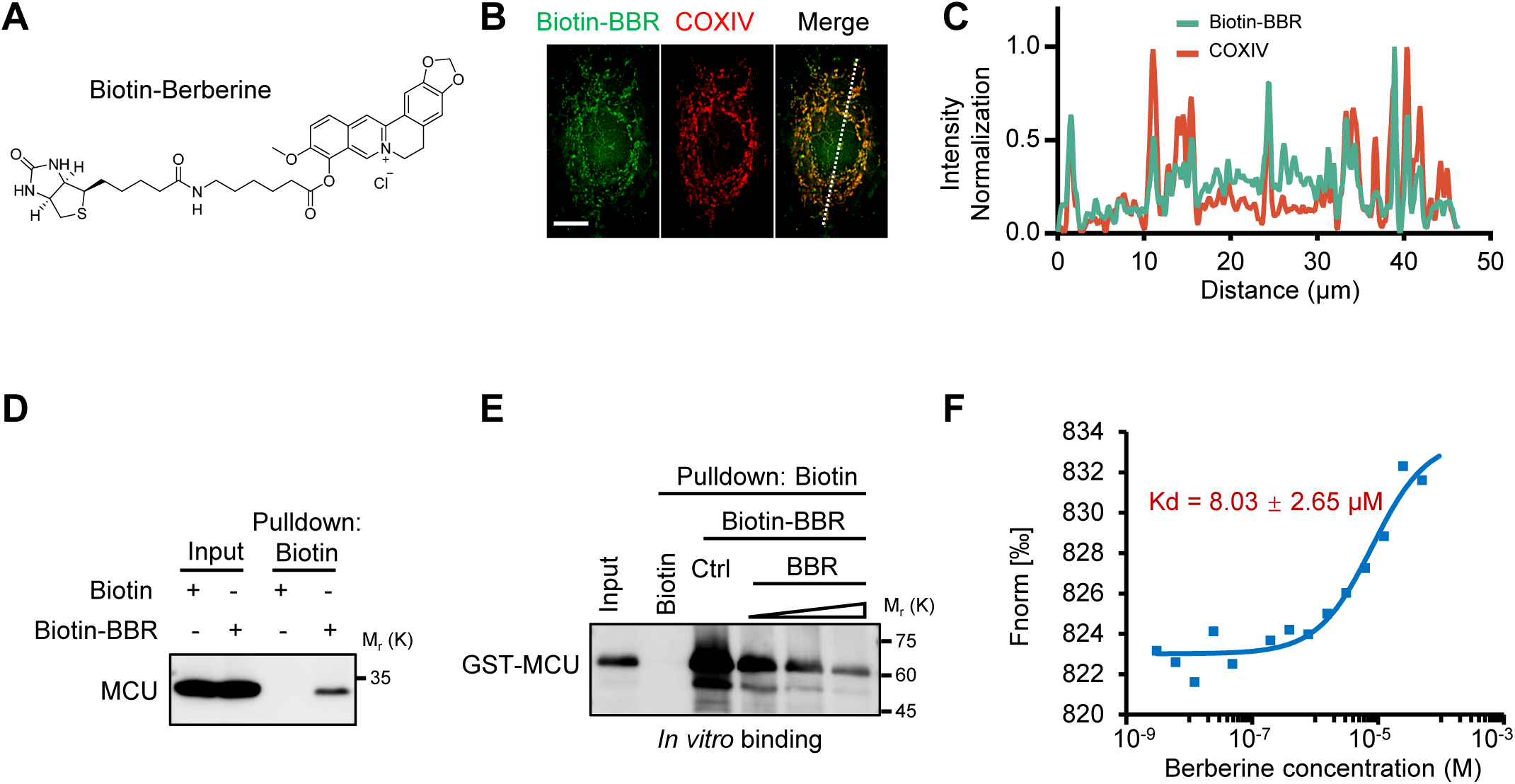
Berberine directly binds to MCU. (A) Chemical structure of Biotin-Berberine. (B) Representative immunofluorescent images of HeLa cells treated with Biotin-Berberine (green) and stained for mitochondria (COXIV, red). Scale bar, 10 μm. (C) Traces of normalized fluorescence intensity spatial profiles through the white line shown in (B). (D) Western blot analysis from Biotin-Berberine pulldown assay using HEK293T cell lysates. Cells were treated with Biotin-Berberine (10 μM, 2 h). (E) Western Blot analysis from *in vitro* pulldown assay investigating interactions between GST-MCU and Biotin-Berberine under varying concentrations of Berberine. (F) Binding ability of Berberine to GST-MCU in MST assays.

**Fig. 3.**
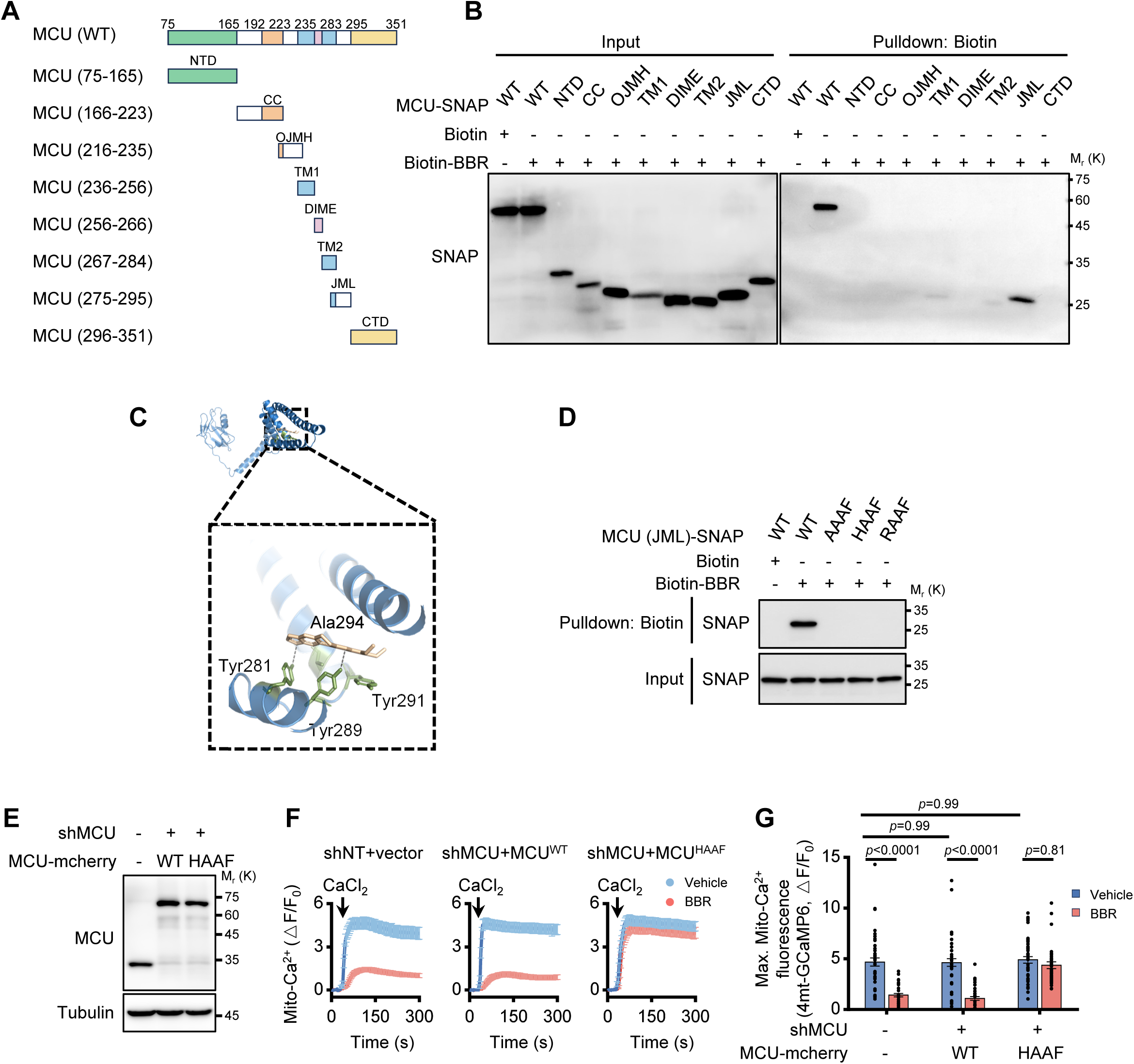
Berberine targets JML domain of MCU to inhibit Ca^2+^ uptake. (A) Schematic representation of MCU truncation mutants utilized for domain-mapping studies. Numerical values in parentheses correspond to the amino acid sequences encompassed by each construct. Key structural domains are labeled: NTD (N-terminal domain), CC (coiled-coil domain), OJMH (outer juxtamembrane helix), TM (transmembrane helix), DIME (DIME motif, Ca^2+^ filter), JML (juxtamembrane loop), and CTD (C-terminal domain). (B) Western blot analysis of Biotin-Berberine pulldown assays using *in vitro* transcribed and translated SNAP-tagged MCU truncation mutants as shown in (A). (C) Molecular docking model revealing the interaction between human MCU and Berberine. (D) Western blot analysis of biotin-berberine pulldown assays using *in vitro* transcribed and translated SNAP-tagged MCU mutants (amino acids 275-295, encompassing the JML region). Wild-type residues Y281, Y289, Y291, and A294 were mutated to AAAF, HAAF, and RAAF, respectively. (E) Western blot analysis of endogenous MCU and exogenous MCU-mCherry expression in HeLa stable cell lines with MCU wild-type (WT) and MCU^HAAF^ mutants. (F) Traces of mitochondrial Ca^2+^ levels, as indicated by the fluorescence intensity of 4mt-GCaMP6, in digitonin-permeabilized HeLa cells following Ca^2+^ addition. The cells include control HeLa, and stable cell lines expressing mcherry-tagged MCU^WT^and MCU^HAAF^ mutants, under conditions with and without 10 μM Berberine. (G) Measurement of the maximal amplitudes of the mitochondrial Ca^2+^ traces in (F). Data are shown as the mean ± s.e.m. n = 45 cells for each condition. *P* value was analyzed using two-way ANOVA.

### Berberine targets the juxtamembrane loop domain of MCU

Next, to explore the mechanism of Berberine’s binding to MCU and its functional impact, we first mapped MCU regions interacting with Berberine. We *in vitro* translated MCU truncations based on its secondary structure (*18*), excluding the mitochondrial targeting sequence (residues 1-74) due to its non-conservative nature (*19*). We assessed their interactions with Berberine using Biotin pull-down assays. Berberine showed a strong affinity for the MCU juxtamembrane loop (JML) segment spanning residues 275-295 (Fig. 3, A and B), although minor binding to other areas was also observed (Fig. 3B). These observations were consistent with molecular docking simulations (Fig. 3C), which indicated that Berberine binds near JML domain, a region critical for EMRE-dependent MCU opening (*18, 20*). Docking analyses pinpointed key binding residues, including Tyr 281, Tyr 289, Tyr 291, and Ala 294 (Fig. 3C, and fig. S3F). Notably, Tyr 281 and Tyr 289 are likely pivotal for hydrogen bonds with Berberine as shown by molecular docking simulations (Fig. 3C, and fig. S3F).

To explore the interaction of Berberine with predicted binding sites within the MCU, we performed site-directed mutagenesis. Sequence alignment of the JML domain revealed four conserved residues (Y281, Y289, Y291, and A294) across various MCU homologs (fig. S3G), known to play a role in maintaining the open conformation of the MCU (*20*). It was reported that mutations of Y289 and Y291 to alanine, as well as A294 to phenylalanine, do not alter the open conformation of MCU and preserve robust mitochondrial Ca^2+^ uptake capacity. However, the critical residue Y281, when mutated to alanine, induces a closed conformation of the MCU (*20*). Under these circumstances, simultaneous mutations of Y281A, Y289A, Y291A, and A294F significantly disrupted the interaction with the Berberine-MCU JML domain (Fig. 3D), confirming Berberine’s direct binding to these residues. To determine whether the ability of Berberine to bind was independent of the MCU’s conformational state (open or closed), we introduced mutations Y281H and Y281R, which maintain the open conformation (*20*) but disrupt potential hydrogen bonding with Berberine. These mutations still abolished Berberine binding (Fig. 3D), negating the hypothesis that Berberine’s binding is confined to the open conformation of MCU.

To further demonstrate that Berberine’s inhibition of the calcium ion absorption function of the MCU is dependent on its binding to the JML domain, we engineered cell lines with stable reintroduction of either the wild-type MCU (MCU^WT^) or the MCU^HAAF^ mutant (Fig. 3E). This experimental design enabled us to contrast the effects of Berberine on mitochondrial Ca^2+^ uptake across these scenarios. Berberine treatment led to a reduction in mitochondrial Ca^2+^ uptake in both control and MCU^WT^-reconstituted cells (Fig. 3, F and G). In contrast, cells expressing the MCU^HAAF^ mutant, which does not bind Berberine due to alterations in the JML domain (Fig. 3D), exhibited no decrease in mitochondrial Ca^2+^ uptake following Berberine treatment (Fig. 3, F and G). This divergence underscores the critical role of the JML domain interaction in Berberine’s mechanism of action on mitochondrial Ca^2+^ regulation.

### Berberine disrupts MCU-EMRE interaction

Ca^2+^ transfer through the mitochondrial inner membrane is mediated by the mitochondrial calcium uniporter (MCU) complex, consisting of MCU, EMRE, MICU1, and MICU2 (*3*). We investigated Berberine’s influence on the MCU complex assembly via co-immunoprecipitation (IP) experiments. The components of MCU complex, including MCU, EMRE, MICU1 and MICU2, were SNAP-tagged and co-expressed with Flag-tagged MCU in 293T cells. Following Berberine treatment, only the interaction between SNAP-tagged EMRE and Flag-tagged MCU was significantly diminished, while the interactions of MCU with other MCU components remained unaltered (Fig. 4A). These results suggest that Berberine selectively disrupts the EMRE-MCU interaction, which was further confirmed by a reverse IP assay (fig. S4A).

**Fig. 4.**
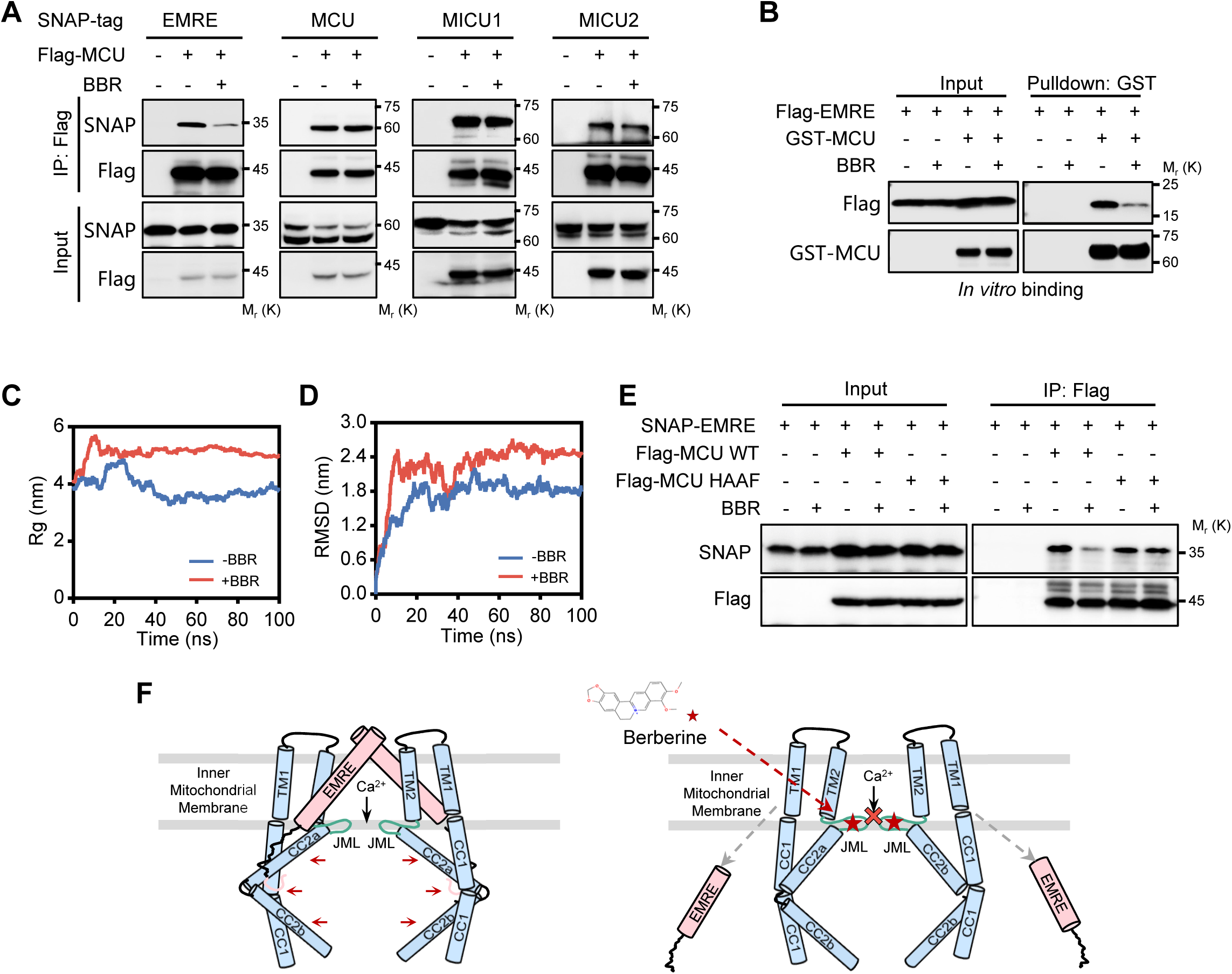
Berberine inhibits Ca^2+^ uptake by disrupting MCU-EMRE complex assembly. (A) Western blot analysis depicting the co-immunoprecipitation of Flag-tagged MCU with SNAP-tagged EMRE, MCU, MICU1, and MICU2 in HEK293T cells, conducted with or without the presence of 10 μM Berberine for 2 hours. (B) Western blot analysis of GST pulldown assays performed with *in vitro* transcribed and translated Flag-tagged EMRE and recombinant GST-MCU, with and without the addition of 2 μg Berberine. (C) Molecular dynamics simulations showing the radius of gyration (Rg) of the MCU-EMRE complex in the presence and absence of Berberine. (D) Molecular dynamics simulations depicting the root-mean-square deviation (RMSD) of the MCU-EMRE complex with and without Berberine. (E) Western blot analysis of the co-precipitation of Flag-tagged MCU wild-type (WT) or Flag-tagged MCU^HAAF^ mutant with SNAP-EMRE in HEK293T cells, treated with 10 μM Berberine for 2 hours or untreated. (F) Proposed model for Berberine-mediated MCU gating. Left panel, EMRE binds to MCU and stabilizes the luminal gate of MCU in the open conformation. Right panel, Berberine integrates into the MCU, inducing a closed conformation that disrupts the assembly of the MCU-EMRE complex and seals the MCU’s luminal gate.

To directly assess Berberine’s effect on MCU-EMRE interaction, we employed *in vitro* binding assays. These assays confirmed Berberine’s ability to directly interfere with MCU-EMRE assembly (Fig. 4B). To understand the mechanism behind this disruption, we conducted molecular dynamics simulations. The simulations revealed that the MCU-EMRE complex rapidly reached equilibrium and maintained stability during the simulation in the absence of Berberine. At 100 ns, the Rg value, a measure of compactness, significantly increased from 3.734 nm to 4.982 nm upon Berberine addition (Fig. 4C), indicating a more spread-out and less compact complex structure. This observation was further corroborated by the increased RMSD values (reflecting average atomic displacement) in the presence of Berberine (Fig. 4D). Together, these results suggest that Berberine induces a looser and less stable complex structure (fig. S4B). Furthermore, hydrogen bond analysis revealed a reduction in the average number of bonds between MCU and EMRE from 10.540 to 6.287 upon Berberine treatment (fig. S4, C and D), signifying a weakening of interactions between the two proteins. This finding aligns with the significant decrease in binding energy observed between MCU and EMRE in the presence of Berberine (from −100.02 ± 17.85 kcal/mol to −29.52 ± 27.07 kcal/mol, fig. S4E). Collectively, these data strongly support the conclusion that Berberine disrupts the MCU-EMRE complex by weakening the interactions between MCU and EMRE.

Next, we sought to clarify that whether Berberine’s disruption of the MCU-EMRE binding activity depends on its interaction with the MCU JML domain. We co-transfected cells with MCU^WT^ or the Berberine (BBR)-binding deficient mutant MCU^HAAF^ alongside EMRE. Both MCU^WT^ and MCU^HAAF^ were capable of interacting with EMRE, indicating that the mutation in the JML domain itself does not affect its assembly with EMRE (Fig. 4E). The addition of Berberine disrupted the assembly between MCU^WT^ and EMRE; however, when Berberine could not bind to the MCU JML domain, as with MCU^HAAF^, Berberine’s presence did not influence the MCU-EMRE assembly. These findings, together with our molecular dynamics simulation results (Fig. 4, C and D, and fig. S4, B to E), suggest that Berberine’s modulation of mitochondrial Ca^2+^ uptake is contingent upon its interaction with MCU. This interaction subsequently disrupts the MCU-EMRE complex assembly and ultimately leads to the inhibition of the MCU channel, thereby preventing Ca^2+^ entry into the mitochondria (Fig. 4F).

### Berberine suppresses mitochondrial Ca^2+^ overload and protects against myocardial ischemia-reperfusion injury

We next sought to explore the physiological relevance of our findings. It’s well established that excessive Ca^2+^ entry into mitochondria can lead to mitochondrial Ca^2+^ overload. This overload subsequently triggers the opening of the permeability transition pore (PTP), causing mitochondrial swelling, and eventually, cell death (*21*). Cardiac-specific conditional knockout of MCU could protect mitochondria by preventing the opening of mPTP *in vitro* and *in vivo* (*10, 11*). The mPTP opening can be monitored by measuring mitochondrial absorbance (*22*). In an *in vitro* context, we observed that Berberine inhibited Ca^2+^ overload-induced mitochondrial swelling (Fig. 5A). This effect paralleled those of Ru360 and cyclosporin A (CsA) (*23*), a known inhibitor of mPTP opening, suggesting that Berberine might confer cellular protection under conditions of mitochondrial Ca^2+^ overload.

**Fig. 5.**
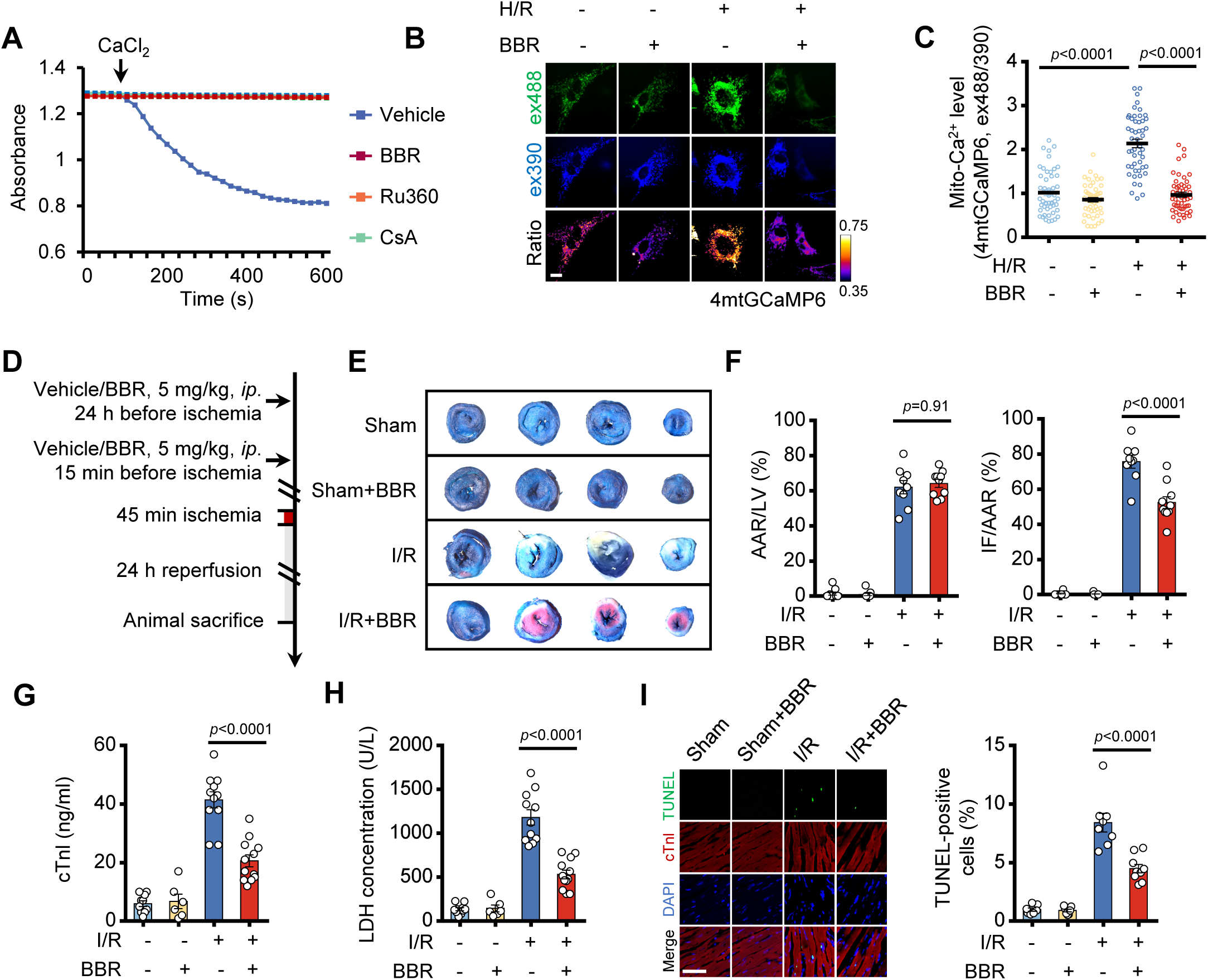
Berberine protects against myocardial ischemia/reperfusion injury through inhibiting mitochondrial Ca^2+^ overload. (A) Mitochondrial swelling in response to Ca^2+^ challenge (500 μM CaCl_2_). Swelling is inhibited with Berberine (10 μM), Ru360 (3 μM) or CsA (3 μM). (B) Representative images of neonatal rat cardiomyocytes expressing 4mtGCaMP6. Cells were either treated with 10 μM Berberine or not, then subjected to hypoxia (3 h) followed by reoxygenation (12 h) (H/R). Fluorescence at ex488 (green) represents the Ca^2+^-dependent signal, while fluorescence at ex390 (blue) indicates the Ca^2+^-independent signal. The ratio (pseudocolour) of ex488/ex390 indicates the level of mitochondrial Ca^2+^. Scale bar, 10 μm. (C) Measurement of mitochondrial Ca^2+^ level indicated by 4mtGCaMP6 fluorescence ratio (ex488/ex405) in (B). Data are shown as the mean ± s.e.m. n = 50 cells for each condition. *P* value was analyzed using one-way ANOVA. (D) The experimental scheme of the mice with acute I/R injury (45 minutes ischemia followed by 24 hours reperfusion) with vehicle or Berberine pretreatment (24 hours and 15 minutes before the onset of ischemia). (E) Evans Blue and 2, 3, 5-triphenyltetrazolium chloride (TTC) staining of myocardium after I/R injury. The blue area indicates non-ischemic area. The red indicates viable tissue at risk. The white indicates infarct area. (F) Quantification of infarct size (IF), area at risk (AAR) of mice subjected to acute I/R injury (as protocols in D). n = 9 or 10 mice. (G-I). Cardiac cell death indexed by serum cTnI (G, n=12 [sham], 6 [sham+BBR], 11 [I/R], and 12 [I/R+BBR]), serum LDH concentration (H, n=13 [sham], 6 [sham+BBR], 11 [I/R], and 12 [I/R+BBR]), and myocardial TUNEL-positive cells (I, n=8 [sham], 6 [sham+BBR], 11 [I/R], and 12 [I/R+BBR]) of the mice subjected to I/R injury (as the protocols in D) with vehicle or Berberine treatment (5 mg/kg, ip.24 h and 15min before I/R injury). *P* value was analyzed using one-way ANOVA. Scale bar, 50 μm.

To validate this hypothesis, we first established a model of hypoxia/reoxygenation (H/R) injury in neonatal rat cardiomyocytes and embryonic rat cardiomyocyte-derived H9c2 cell line. Consistent with previous studies (*24, 25*), H/R injury led to a significantly elevated mitochondrial Ca^2+^ levels, indicated by fluorescent intensity of 4mtGCaMP6 or Rhod2 (Fig. 5, B and C, and fig. S5, A and B). Furthermore, Berberine pretreatment alleviated this mitochondrial Ca^2+^ overload (Fig. 5, B and C, and fig. S5, A and B). To assess the potential cardioprotective effect of Berberine *in vivo*, a mouse acute cardiac ischemia/reperfusion (I/R) model was established (Fig. 5D). Pretreatment of Berberine (24h plus 15 min before reperfusion) significantly reduced I/R induced myocardial infarct size (Fig. 5, E and F). Additionally, Berberine ameliorated cardiomyocyte death, evidenced by decreased cTNl and LDH concentration (Fig. 5, G and H). TUNEL staining experiments further confirmed the protective effect of Berberine against myocadiac I/R injury (Fig. 5I). In summary, these data indicate that Berberine exhibits therapeutic effects in myocardial ischemia/reperfusion injury via counteracting mitochondrial Ca^2+^ overload.

## Discussion

Our virtual screening approach using an FDA-approved drug library identified Berberine, a natural alkaloid derived from medicinal plants, as a potent MCU inhibitor. Berberine exerts its inhibitory effect by directly binding to the MCU and disrupting the assembly of MCU-EMRE complex, consequently hindering Ca^2+^ entry into mitochondrial matrix. Notably, Berberine significantly reduces mitochondrial Ca^2+^ overload, thereby protecting cardiomyocytes from ischemia-reperfusion (I/R)-induced apoptosis *in vitro* and *in vivo*. These results underscore the potential of Berberine as a promising MCU-inhibiting drug for clinical application.

Berberine, a well-characterized plant alkaloid with a long history of traditional use, has been employed for treating various conditions, including diabetes, hyperlipidemia, and cardiovascular diseases (*26*). Its antimicrobial properties are also well-documented (*27*). Despite its extensive therapeutic application, Berberine’s precise molecular targets remains elusive. Our study provides compelling evidence that Berberine targets MCU, effectively impeding mitochondrial Ca^2+^ uptake through direct binding. Importantly, Berberine may overcome limitations observed in existing MCU inhibitors such as Ru360, DS16570511 (*8*), mitoxantrone (*9*), and MCU-i11 (*28*)—a specific MICU1-targeting inhibitor—in animal experiments. DS16570511, for instance, exhibits off-target effects on mitochondrial membrane potential, thereby limiting its application (*29*). Similarly, mitoxantrone, a topoisomerase inhibitor, is known to have cardiotoxic effects (*30*). Berberine’s well-established safety profile *in vivo* positions it as a strong candidate for further exploration in clinical settings.

Our studies, along with previous studies, consistently demonstrate Berberine’s capacity to provide significant cardioprotection in the context of myocardial I/R injury (*31*). Clinical studies further support its potential, indicating that Berberine can modulate inflammatory markers in patients suffering from acute myocardial infarction (*32*). Berberine’s protective effects are multifaceted, encompassing antioxidant, anti-inflammatory, and anti-apoptotic activities. A key aspect of this protection is Berberine’s ability to maintain mitochondrial integrity and high activity of the respiratory complexes (*33–35*), as evidenced by its regulation of Bcl-2 and Bax proteins, which are crucial for mitochondrial membrane stability and the prevention of apoptotic factor release (*33, 36*). Our findings establish MCU as a key cardiac target of Berberine, revealing a core mechanism for its protective effects in I/R injury. Given that mitochondrial Ca^2+^ overload occurring rapidly within the initial 20 minutes post-ischemia (*25*), early intervention with Berberine is of paramount importance. Furthermore, Berberine’s potential interactions with other targets and its slower-acting mechanisms warrant further investigation to fully understand its therapeutic potential.

Our findings reveal that Berberine binds directly to the MCU near the juxtamembrane loop (JML) domain, at residues Y281, Y289, Y291, and A294. The JML is essential for channel gate formation and its interaction with EMRE is critical for Ca^2+^ channel activation (*18*). Mutational analysis of Y281 and Y289 highlights their significance in MCU-EMRE assembly and the EMRE-mediated conformational opening of MCU (*20*). Y281A mutants significantly disrupts EMRE binding (*20*), paralleling the loss of Ca^2+^ uptake observed with Berberine treatment. Notably, Y289, a key hydrophobic residue within the JML, facilitates EMRE-dependent MCU opening (*20*). Substituting Y289 with hydrophilic residues impairs channel opening, underscoring the critical role of hydrophobic interactions at this site for EMRE binding and MCU activation. Berberine’s hydrophobic surface (*37*) might promote its integration near the JML, disrupting interactions between critical residues like Y281 and Y289, and fostering a closed MCU conformation unfavorable to EMRE association. While our molecular dynamics simulations support this hypothesis, the complex mechanisms of MCU gating warrant further structural studies for full elucidation.

In conclusion, our study identifies Berberine as a novel MCU inhibitor with significant potential for clinical applications. Revealing its inhibitory effects on MCU and providing insights into the complex Berberine-MCU-EMRE dynamics, our findings lay a solid foundation for the development of targeted drug strategies aimed at mitochondrial Ca^2+^ regulation. Structural analogs of Berberine could be engineered for enhanced specificity and safety as MCU inhibitors. Furthermore, the role of MCU dysfunction is becoming increasingly understood, not only in cardiac ischemia/reperfusion injury but also for other diseases, such as the recently reported dependency of venetoclax-resistant leukemic stem cells (LSCs) on mitochondrial calcium for survival(*38*). These findings position Berberine as a valuable starting point for developing novel interventions in diseases associated with dysregulated mitochondrial Ca^2+^ homeostasis.

## Materials and Methods

### Experimental animals

All animals used in this study were maintained in the pathogen-free barrier animal facility at the National Center of Biomedical Analysis. Animal care was monitored daily by certified veterinary staff and laboratory personnel. 10-week-old mice in the C57BL/6J background and pregnant Sprague-Dawley rats were purchased from the Beijing Vital River Laboratory Animal Technology Company.

### Murine model of myocardial ischemia

The 10 week-male-C57BL/6 mice (>22 g) were randomly assigned to specific groups. Berberine (5 mg/kg) or vehicle was administered intraperitoneally twice (24 hours and 15 minutes prior) to ischemia-reperfusion (I/R) surgery as described (*39*). Briefly, pentobarbital (70 mg/kg i.p.) was used to induce and maintain anesthesia. Mice were then ventilated on a Harvard rodent respirator via a tracheostomy. A midline sternotomy was performed, with a reversible snare occlude placed around the left anterior descending coronary artery. I/R injury was initiated by tightening the snare for 45 minutes, then releasing it. After reperfusion for 24 hours, the mice were sacrificed for further detection and measurement.

For serum LDH concentration and cTnl level, blood samples were centrifuged at 3,000 rpm for 10 min and then detected using an LDH kit (Sigma-Aldrich, Cat# MAK066) and a mouse cardiac troponin I ELISA kit (ThermoFisher, Cat#EEL112) according to the manufacturer’s instructions.

For the measurement of the area-at-risk (AAR) and infarct size, the heart was initially excised post-reocclusion of the coronary artery at the previous occlusion site. The heart ascending aorta was cannulated (distal to the sinus of Valsalva) and then perfused retrogradely with 0.05% Alcian blue to delineate the AAR. Following AAR visualization, the heart was frozen at −80 L for 5 min, cut into ∼1 mm slices, and the unstained infarcted (IF) region was lucifugal visualized using 1% 2,3,5-triphenyl-tetrazolium chloride at 37 L for 15 min. Infarct and left ventricular areas were determined by planimetry with Image J. The infarct size was calculated as infarct area divided by area at risk (IF/AAR).

### Molecular Docking Screening

The 3D structures of MCU (PDB: 6XJV) were retrieved from the Protein Data Bank. A set of 2,816 FDA-approved drugs were converted to 3D structures using Open Babel. Preparation and parameterization of the receptor protein and ligands were conducted with AutoDock Tools (ADT3). Docking grid documents were created using AutoGrid of sitemap, and docking simulation were performed with AutoDock Vina (1.2.0) (*40*). The score was ranked according to the value of the binding energy of protein-ligand complex. Finally, the protein-ligand interaction figure was generated by PyMOL.

### Cell culture

To establish HeLa cells stably expressing 4mt-GCaMP6s, we constructed a lentiviral vector (pLVX-puro) containing the 4mt-GCaMP6s insert. This vector was transfected into HEK293T cells along with the packaging plasmids pSPAX2 and pVSV-G using a 4:3:1 ratio. Stable transfectants were selected using 1 μg/ml puromycin. HeLa cells were cultured at 37°C in DMEM supplemented with 10% FBS and 1% penicillin-streptomycin. A similar approach was employed to express MCU WT and MCU HAAF in the HeLa cell line. Initially, HeLa cells were infected with the pLKO.1-shMCU lentivirus and subsequently selected with 1 μg/ml puromycin. Knockout HeLa cells were then infected with lentivirus carrying pLVX-mCherry-N1 vectors harboring either MCU^WT^ or MCU^HAAF^. Cells expressing mCherry were isolated using flow cytometry. Knockout of endogenous MCU and expression of exogenous MCU-mCherry were confirmed by Western blot analysis. All other mammalian cell lines used in this study were from the American Type Culture Collection. All the cell lines were fully authenticated and tested free of mycoplasma.

### Mitochondrial Ca^2+^ modulator Screening

Hela cells expressing 4mt-GCaMP6s were permeabilized by a 1 min perfusion with intracellular buffer (130 mM KCl, 10 mM NaCl, 2 mM K_2_HPO_4_, 5 mM Succinic acid, 5 mM Malic acid, 1 mM Pyruvate, 0.5 mM ATP, 0.1 mM ADP, MgCl_2_, 20 mM HEPES, pH 7, 37 L) supplemented with 10 μM digitonin and 50 μM EGTA. Then the buffer was changed to intracellular buffer supplemented with 10 μM compounds from primary screen hits of FDA-approved drugs for 10 min. Mitochondrial Ca^2+^ uptake was evoked by adding 100 μM CaCl_2_ to the buffer. Ca^2+^ imaging was performed using a DeltaVision Deconvolution microscope (GE Healthcare) equipped with a ×20 objective. Exposure time was set to 25 ms for FITC. Images were acquired at 1 frame per 3 seconds with an EM-CCD camera. Images were analysed using Image J software. All images were background-corrected by subtracting mean pixel values of a cell-free region of interest.

### Molecular dynamics (MD) simulations

Structural data for MCU and EMRE monomers were retrieved from the Protein Data Bank (PDB) via the entry code 6XJV. The preprocessing phase employed Gromacs 2018 as the simulation suite, coupled with the Amber99SB-ildn force field for encompassing protein and small molecule interactions. A 10×10×10 nm^3^ water box, utilizing the TIP3P model, was constructed to solvate the system, ensuring a minimal 1.2 nm buffer between the protein surfaces and the box boundaries. This was followed by ion addition for electrostatic equilibrium.

For treating electrostatic interactions, we implemented the Particle-mesh Ewald (PME) method. Energy minimization was executed through the steepest descent method over 50,000 steps, setting both the Coulombic and van der Waals interaction cutoffs at 1 nm. Equilibration phases under NVT (constant number of particles, volume, and temperature) and NPT (constant number of of particles, pressure, and temperature) settings preceded a 100 ns MD simulation, conducted under standard laboratory conditions of temperature and pressure. Non-bonded interactions were applied with a 10 Å cutoff. Thermal and pressure stability were maintained at 300 K and 1 bar, respectively, utilizing the V-rescale thermostat and the Berendsen barostat.

This rigorous preparation and execution strategy yielded a robust framework for accurately simulating the complex’s behavior in a physiological context, culminating in the acquisition of data such as radius of gyration (Rg), root-mean-square deviation (RMSD), number of hydrogen bonds (H-bonds), and total free energy, to assess structural stability, conformational dynamics, interaction strength, and energetics throughout the simulation period.

### Ca^2+^ measurements

For imaging Ca^2+^ transients evoked by treatments with an agonist, cells labelled with Calbryte590 or 4mt-GCaMP6 were incubated at 37 L with Krebs-Ringer modified buffer (KRB), which contains 125 mM NaCl, 5 mM KCl, 1 mM Na_3_PO_4_, 1 mM MgSO_4_, 5.5 mM glucose and 20 mM HEPES (pH 7.4). After 10 μM histamine stimulation, a series of images were captured every 3 s using a DeltaVision Deconvolution microscope (GE Healthcare). The recorded images were analyzed and quantified using Image J software (NIH).

For image analysis, background correction was first performed frame by frame by subtracting the intensity of a nearby cell-free region from the signal of the imaged cell. F_0_ is calculated as the initial background-subtracted fluorescence intensity. F indicates the background-subtracted fluorescence intensity at each time point. ΔF/F_0_= (F – F_0_)/F_0_, indicates the Ca^2+^ concentration change after stimulation.

### Mitochondrial membrane potential measurements

For confocal microscopy, adherent HeLa cells were stained with Tetramethylrhodamine (TMRM) at a final concentration of 100 nM in fresh culture medium. After a 30-minute incubation period at 37°C to allow for adequate dye uptake, cells were treated with the compounds of interest for an additional 30 minutes at the same temperature. Subsequently, cells were washed to remove excess dye and unbound compounds. Imaging was performed on a DeltaVision confocal microscope (GE Healthcare). Acquired images were then processed and quantified using ImageJ software.

For flow cytometric analysis, Hela cells were digested and stained with 100 nM Tetramethylrhodamine (TMRM) in fresh medium for 30 min at 37L. Compounds were added for 30 min at 37L. Then cells were washed twice and resuspended in PBS containing 7AAD. Samples were analyzed using cytoFLEX flow cytometer (Beckman Coulter).

### Western blot analysis

For western blot analysis, cells were lysed in RIPA buffer (50 mM Tris-HCl (pH 8.0), 150 mM NaCl, 1% (v/v) Nonidet P-40, 0.5% sodium deoxycholate, 0.1% SDS, protease inhibitor cocktail). Protein samples were resolved by SDS-PAGE and transferred onto PVDF membranes. Membranes were immunoblotted with the indicated primary antibodies. Antibodies were used at the following concentrations: anti-MCU, rabbit polyclonal, Cell Signaling Technology, Cat# 14997 (1:1000); anti-SNAP, rabbit polyclonal, New England Biolabs, Cat# P9310S (1:1000).

### Immunoprecipitation

HEK293T cells were transfected with the indicated plasmids using Lipofectamine^TM^ 2000 transfection reagent (ThermoFisher, Cat# 11668500). After transfection for 24 h, cells were treated with 10 μM Berberine for 2 hours before harvested. Cells were lysed with RIPA buffer (50 mM Tris-HCl (pH 8.0), 150 mM NaCl, 1% (v/v) Nonidet P-40, 0.5% sodium deoxycholate, protease inhibitor cocktail).

Cell lysates were immunoprecipitated by anti-Flag M2 affinity gel (Selleck, Cat# B23102) overnight at 4 L. Beads were washed 5 times in RIPA buffer followed by eluting bound protein complexes in 2× SDS dye. Whole cell lysates and co-precipitation samples were analyzed by western blot.

### Biotin pulldown assay

For *in vivo* Biotin pulldown assay, HEK293T cells were treated with Biotin or Biotin-labeled Berberine (10 μM, 2 h). Cells were lysed and harvested with RIPA buffer, which containing 50 mM Tris-HCl, 150 mM NaCl, 1% NP-40 and 0.5% sodium deoxycholate. Balanced Streptavidin Agarose beads (ThermoFisher, Cat# 20357) were added and incubated overnight at 4 °C. The beads were washed with RIPA buffer adding 0.03% Tween 5 minutes for 5 times. The indicated proteins were detected by Immunoblotting.

For the *in vitro* biotin pulldown assay, biotin or biotin-labeled berberine (2 μg) was pre-incubated with either recombinant protein (2 μg) or in vitro transcribed/translated protein (Promega, Cat# L1170) for 8 hours at 4°C. The mixture was then incubated with streptavidin agarose beads (Thermo Fisher, Cat# 20357) overnight at 4°C. The beads were washed as previously described and the bound proteins were analyzed by immunoblotting.

### Immunofluorescence and confocal microscopy

For visualizing the intracellular distribution of Berberine, HeLa cells were grown on coverslips, incubated with 50 nM MitoTracker^TM^ DeepRed (ThermoFisher, Cat# M22426) in fresh medium for 15 min at 37 °C. 10 μM Berberine was added for 10 min. After treatment, the medium was replaced with phenol red-free DMEM supplemental with 10% FBS. Then cells were visualized using a DeltaVision Deconvolution microscope (GE Healthcare). Images of Berberine and MitoTracker^TM^ DeepRed signals, acquired by FITC and Cy5 filters respectively, were merged to visualize their co-localization in mitochondria.

For visualizing the intracellular distribution of Biotin-labeled Berberine, HeLa cells were grown on coverslips, incubated with 10 μM Biotin-Berberine for 10 min at 37 °C. After treatment, cells were fixed in 4% paraformaldehyde for 5 min. Next, the cells were incubated with 0.3% Triton X-100 in PBS for 10 min on ice and then blocked in 3% BSA in PBS for at least 1 h. The primary antibody was incubated overnight at 4°C followed by secondary antibody incubation. Finally, the cells were mounted using 80% glycerol. Antibodies were used at the following concentrations: anti-COXIV, rabbit polyclonal, Cell Signaling Technology, Cat# 4850 (1:200); Alexa Fluor 488-conjugated streptavidin antibody, ThermoFisher, Cat# S11223 (1:400); Alexa Fluor 546-conjugated anti-Rabbit Secondary Antibody, ThermoFisher, Cat# A11035 (1:400). Images were acquired using a DeltaVision Deconvolution microscope.

### Mitochondrial isolation and swelling assay

For isolating cardiac mitochondria, mouse hearts were minced in mitochondrial isotonic buffer (225 mM mannitol, 75 mM sucrose, 5 mM MOPS, 0.5 mM EGTA and 2 mM taurine, pH 7.25) supplemented with protease inhibitor cocktail, and subsequently homogenized using a dounce homogenizer. The homogenate was initially centrifuged for 10 min at 700 × g and the resulting supernatant was next spun for 10 min at 12,000 × g to pellet the mitochondria.

Mitochondrial swelling was measured as previously described (*23*). In brief, isolated cardiac mitochondria (100 μg) were resuspended in swelling buffer (120 mM KCl, 10 mM Tris-HCl, 5 mM MOPS, 5 mM Na_2_HPO_4_, 10 mM glutamate, 2 mM malate and 0.1 mM EGTA) in a total volume of 200 μl. Pore opening was induced and detected by the addition of 500 μM of total CaCl_2_ while monitoring absorbance at 540 nm. Where indicated, Berberine (10 μM), cyclosporin A (3 μM) or Ru360 (3 μM) was added.

### Isolation and culture of primary neonatal rate cardiomyocytes (NRCMs)

NRCMs were isolated and cultured as described previously (*41*). Briefly, cardiomyocytes were isolated from the neonatal hearts of <24h-old Sprague-Dawley rats using PBS containing 0.03% trypsin and 0.04% collagenase type II. The digestion was halted by Dulbecco’s modified Eagle’s medium (DMEM), supplemented with 10% fetal bovine serum (FBS). After differential attachment for 90 min, the cells in suspension were then plated and cultured in DMEM with 10% FBS at 37°C with 5% CO_2_ for 24 h.

### Hypoxia/reoxygenation (H/R) injury model

The hypoxia/reoxygenation (H/R) injury of NRCMs or H9c2 cells were achieved in incubator (Thermo Fisher Scientific, Madison, WI, USA) filled with 94 % N_2_, 5 % CO_2_, and 1% O_2_ in DMEM deprived of glucose and sodium pyruvate for 4h, followed by 8 h-reoxygenation by transferring the cells to a normoxia incubator with normal supplement of oxygen and glucose.

To examine the protective effect of BBR on H/R-induced Ca^2+^ overload, NRCMs infected with Ad-4mt-GCaMP6 or H9c2 cells stained with Rhod2 were treated with BBR at a final concentration 10 μM throughout the H/R period. Then cells were visualized using a DeltaVision Deconvolution microscope (GE Healthcare). To assess mitochondrial Ca^2+^ level using Rhod2, cells labeled with Rhod2 were illuminated at 546 nm, and fluorescence was collected through a TRITC filter. To assess mitochondrial Ca^2+^ level using 4mt-GCaMP6, cells labeled with 4mt-GCaMP6 were illuminated at 488 and 390 nm, and fluorescence was collected through a FITC filter. Images were analysed using Image J software. All images were background-corrected by subtracting mean pixel values of a cell-free region of interest.

### Statistics and reproducibility

Statistical comparisons between only two groups were carried out using unpaired Student’s t test or the Mann-Whiney U test when a normal distribution could not be assumed. For comparisons across multiple groups, a one-way analysis of variance (ANOVA) was employed. Furthermore, when examining the effects of two independent variables or the interaction between them on a dependent variable, a two-way ANOVA was utilized to discern any significant differences. Statistical calculations were carried out using GraphPad Prism 6.0. We tested data for normality and variance, and considered a *P* value of less than 0.05 as significant. All experiments were performed three or more times independently under identical or similar conditions.

## Acknowledgments

The authors thank D.D. Stefani for providing 4mt-GCaMP6 plasmids, and K. Wang and X. Xu for assistance with microscopy assays. This work was supported by the National Natural Science Foundation of China (grant numbers 82025028, 92354304 and 82000107), the National Key R&D Program of China (2021YFA1300203 and 2020YFA0113300), China Postdoctoral Science Foundation (2022M723897).

## Author Contributions

Xin Pan conceived the study. Haixin Zhao designed and carried out most of the experiments with the assistance from Siqi Chen, Nian Cao, and Wenjun Wu. Haixin Zhao, Siqi Chen, Guangqin Liu and Haizhen Zhu performed calcium measurements and biochemical experiments. Haixin Zhao, Nian Cao and Jun Gao contributed to mouse studies. Haixin Zhao and Wenjun Wu performed Screening and Data Analysis. Dingyi Lu and Lingmin Zeng performed immunostaining and flow cytometry analysis. Jiayi Chen, Teng Li, Ting Li, Weina Zhang, Qing Xia, Tao Zhou, Ai-Ling Li, Xue-Min Zhang, and Xin Pan contributed to interpreting the results. Xin Pan and Ai-Ling Li supervised the research. Xin Pan and Haixin Zhao wrote the paper.

## Supplementary Material

**Fig. S1. Screening identifies Berberine as an inhibitor of mitochondrial Ca^2+^ uptake.** (A) Histogram plot of the 2,816 FDA-approved drug library based on the free-energy docking score with MCU. (B) Traces of mitochondrial Ca^2+^ level (indicated by fluorescence of 4mt-GCaMP6) in digitonin-permeabilized HeLa cells upon the addition of Ca^2+^ in the presence of DMSO, Ru360, or drugs. (C) Graphical representation of drug screen results. ‘Z-score’ ranks drugs based on maximal amplitudes of the traces in B. (D) The list of the top 7 compounds with mitochondrial Ca^2+^ uptake inhibitory activity. (E) Measurement of mitochondrial membrane potential indicated by TMRM fluorescence intensity. Data are shown as the mean ± s.e.m. n = 50 cells for each condition. *P* value was analyzed using one-way ANOVA.

**Fig. S2. Berberine selectively inhibits mitochondrial Ca^2+^ uptake without impairing cytosolic Ca^2+^ signaling, mitochondrial membrane potential, or cell viability.** (A) Representative traces of cytosolic Ca^2+^ level (indicated by GCaMP6) in 10 μM Histamine-stimulated HeLa cells pretreated with different concentration of Berberine. Data are shown as the mean ± s.e.m. n = 50 cells for each condition. (B) Measurement of the maximal amplitudes of the cytosolic Ca^2+^ traces in A. Data are shown as the mean ± s.e.m. n = 50 cells for each condition. *P* value was analyzed using one-way ANOVA. (C) Representative traces of mitochondrial Ca^2+^ level (indicated by 4mt-GCaMP8) in 10 μM Histamine-stimulated HeLa cells pretreated with different concentration of Berberine. Data are shown as the mean ± s.e.m. n = 50 cells for each condition. (D) Measurement of the maximal amplitudes of the mitochondrial Ca^2+^ traces in C. Data are shown as the mean ± s.e.m. n = 50 cells for each condition. *P* value was analyzed using one-way ANOVA. (E) Flow cytometry analysis of TMRM staining in HeLa cells pretreated with different concentrations of Berberine and 50 μM CCCP. Shown is one experiment that is representative of three similar experiments. (F) Quantitative analysis of relative fluorescence of TMRM staining after different concentrations of Berberine treatment and 50 μM CCCP treatment. Data are shown as the mean ± s.e.m. n = 3 independent experiments. (G) Cell viability of HeLa cells, U2OS cells and HEK293T cells after different concentration of Berberine treatment was assessed at indicated time and normalized to day 0 in each group. Data are shown as the mean ± s.e.m. n = 3.

**Fig. S3. Berberine directly targets MCU.** (A) Representative images of Berberine (left panel) treated HeLa cells co-stained with MitoTracker DeepRed (middle panel). The merged image is shown in the right panel. Scale bar, 10 μm. (B) Traces of normalized fluorescence intensity spatial profiles through the white line shown in A. (C) Traces of mitochondrial Ca^2+^ level (indicated by fluorescence of 4mt-GCaMP6) in digitonin-permeabilized HeLa cells upon the addition of Ca^2+^ in the presence of different concentration of Berberine or Biotin-Berberine. Data are shown as the mean ± s.e.m. n = 40 cells for each condition. (D) Measurement of the maximal amplitudes of the mitochondrial Ca^2+^ traces in C. Data are shown as the mean ± s.e.m. n = 40 cells for each condition. *P* value was analyzed using two-way ANOVA. (E) Western Blot analysis of Biotin-Berberine pulldown assay in MCU-Flag overexpressed HEK293T cells. Cells were treated with 2 h 10 μM Biotin or 10 μM Biotin-Berberine. (F) Two-dimensional diagrams illustrating the binding mode of berberine with MCU. (G) Sequence alignment of the putative region (amino acids 275-295) surrounding the JML domain of MCU proteins from 16 diverse species.

**Fig. S4. Berberine disrupts MCU-EMRE assembly.** (A) Western Blot analysis of co-precipitation of the Flag-EMRE with SNAP-MCU in HEK293T cells in the presence of 10 μM Berberine or not for 2 hours. (B) Molecular dynamics simulations illustrating the assembly conformations of the MCU-EMRE complex in the presence and absence of Berberine. (C) Molecular dynamics simulations reveal the hydrogen bonding interactions between MCU and EMRE under conditions with and without Berberine. (D) Molecular dynamics simulations displaying the number of hydrogen bonds between MCU and EMRE in the presence and absence of Berberine. (E) Molecular dynamics simulations demonstrating the binding free energy between MCU and EMRE with and without Berberine. The components of the binding free energy are detailed as follows: van der Waals energy (VDWAALS), electrostatic energy (EEL), polar solvation energy (EGB), non-polar solvation energy (ESURF), total gas phase free energy (GGAS, calculated as the sum of VDWAALS and EEL), total solvation free energy (GSOLV, calculated as the sum of EGB and ESURF), and the total free energy (TOTAL, calculated as the sum of GSOLV and GGAS).

**Fig. S5. Berberine attenuates mitochondrial Ca^2+^ overload in cardiomyocytes under pathological conditions.** (A-B) Measurement of mitochondrial Ca^2+^ level (indicated by Rhod2) in H9C2 cells. Cells were pretreated with 10 μM Berberine or not, and then were subjected to hypoxia (3 h) followed by reoxygenation (12 h) (H/R). Relative fluorescence intensity of Rhod2 were measured in B. Data are shown as the mean ± s.e.m. n = 30 cells (Control and BBR groups), n = 40 cells (H/R and H/R+BBR groups). *P* value was analyzed using one-way ANOVA. Scale bar, 10 μm.

