## Supplementary Figures for "Berberine is a novel mitochondrial calcium uniporter (MCU) inhibitor that disrupts MCU-EMRE assembly"

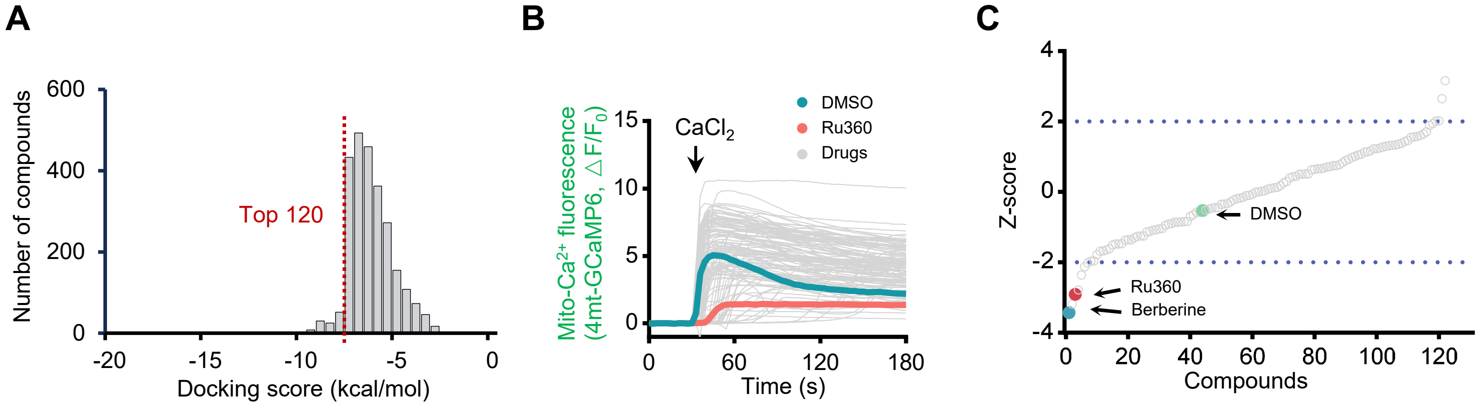


D

| Compound (10 μM) | Inhibition (%) | *P* value |
|---|---|---|
| Berberine | 87.21264 | < 0.0001 |
| Trifarotene | 79.77925 | < 0.0001 |
| Ru360 | 71.36228 | < 0.0001 |
| Eltrombopag | 67.76502 | < 0.0001 |
| Tipranavir | 55.00618 | < 0.0001 |
| Oxyphenisatine | 47.85387 | < 0.0001 |
| Ramatroban | 44.67031 | < 0.0001 |

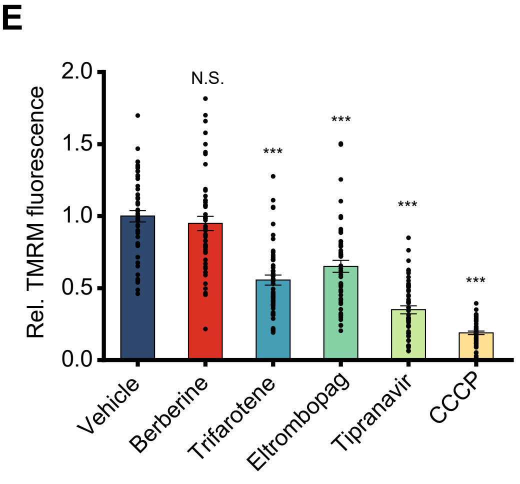


**Supplementary Figure 1. Screening identifies Berberine as an inhibitor of mitochondrial $Ca^{2+}$ uptake.**

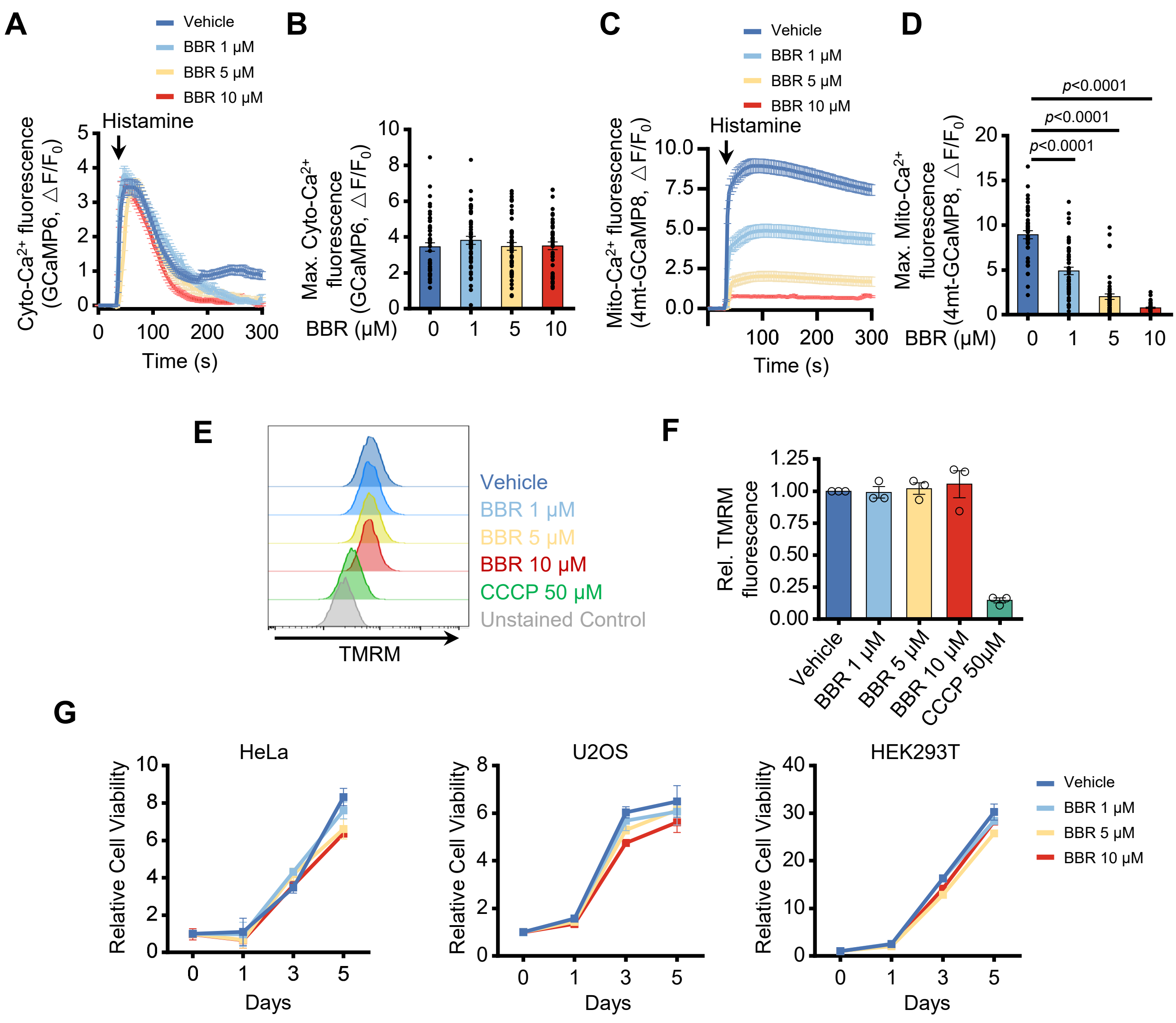


**Supplementary Figure 2. Berberine selectively inhibits mitochondrial $Ca^{2+}$ uptake without impairing cytosolic $Ca^{2+}$ signaling, mitochondrial membrane potential, or cell viability.**

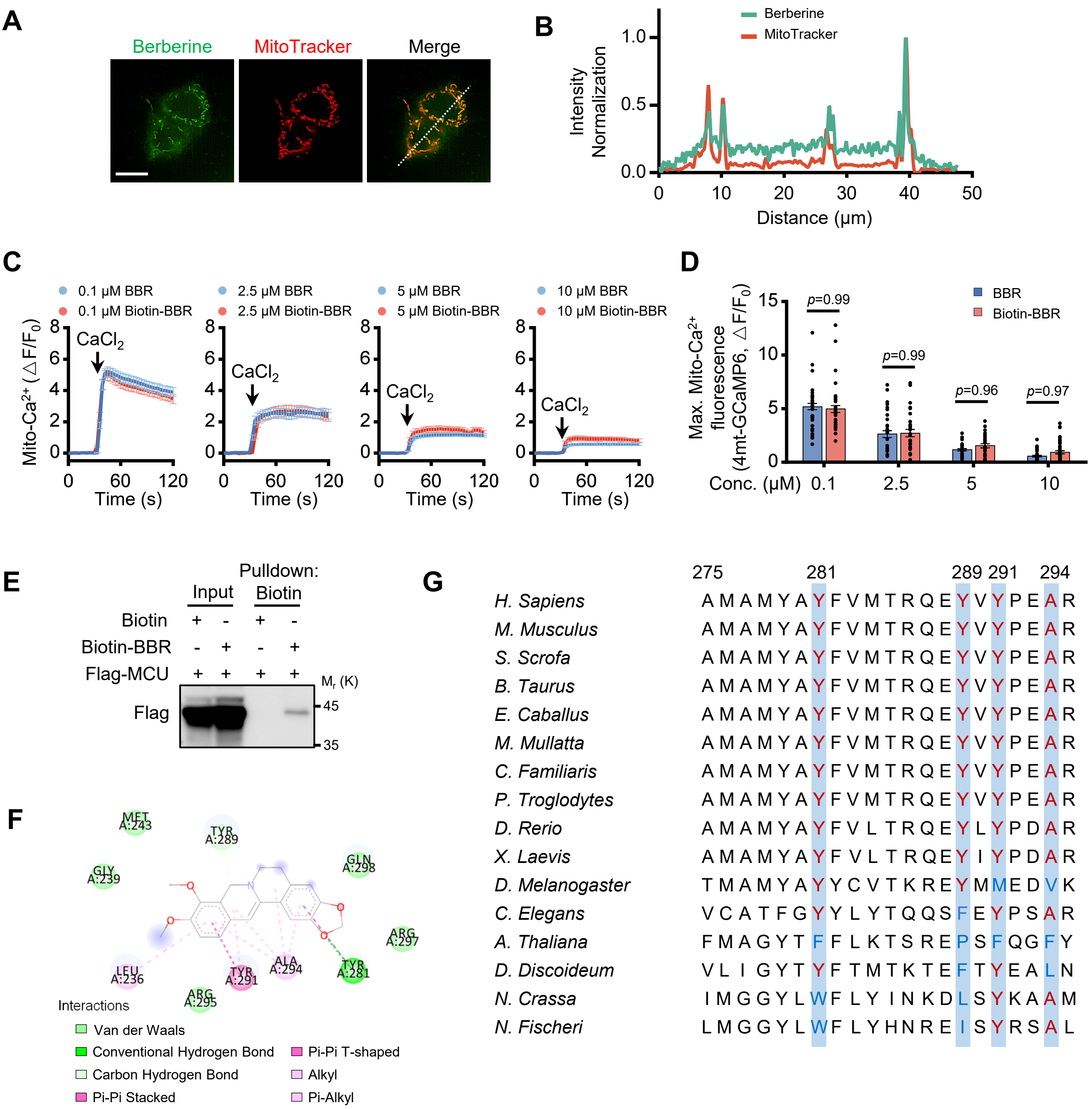


**Supplementary Figure 3. Berberine directly targets MCU.**

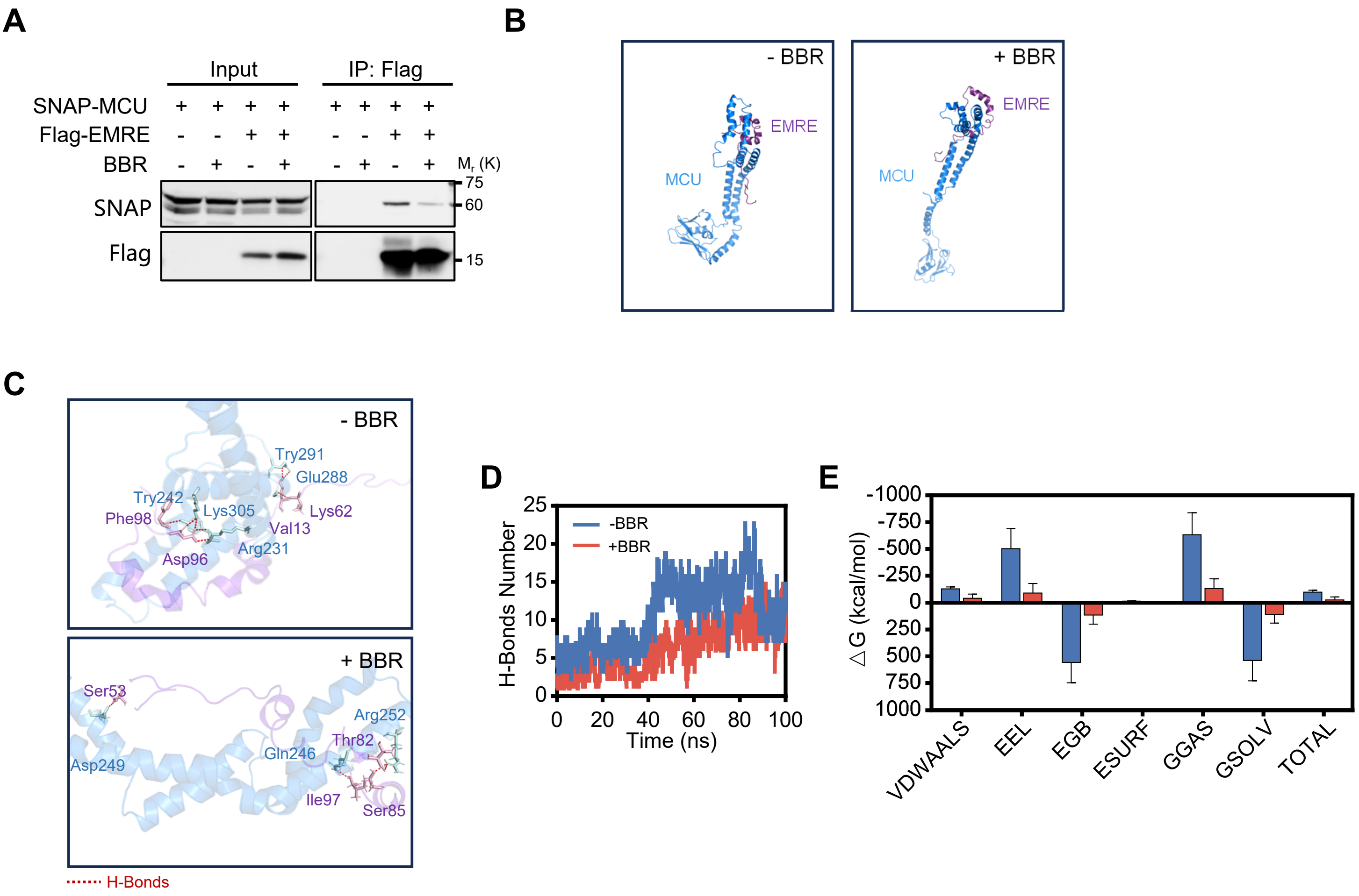


**Supplementary Figure 4. Berberine disrupts MCU-EMRE assembly.**

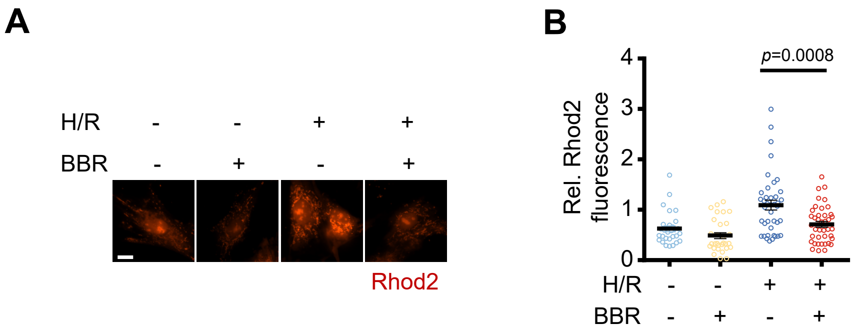


**Supplementary Figure 5. Berberine attenuates mitochondrial $Ca^{2+}$ overload in cardiomyocytes under pathological conditions.**
